# The antagonistic transcription factors, EspM and EspN, regulate the ESX-1 secretion system in *M. marinum*

**DOI:** 10.1101/2024.01.09.574899

**Authors:** Kathleen R. Nicholson, Rachel M. Cronin, Rebecca J. Prest, Aruna R. Menon, Yuwei Yang, Madeleine K. Jennisch, Matthew M. Champion, David M. Tobin, Patricia A. Champion

**Affiliations:** Eck Institute for Global Health, Department of Biological Sciences, University of Notre Dame, Notre Dame, Indiana, USA; Department of Molecular Genetics and Microbiology; Department of Immunology, Duke University School of Medicine, Durham, North Carolina USA; Department of Chemistry and Biochemistry, University of Notre Dame, Notre Dame, Indiana, USA; Department of Microbiology-Immunology, Northwestern University Feinberg School of Medicine, Chicago, Illinois, USA

**Keywords:** Mycobacterium, Type VII Secretion, ESX-1, Regulation, Transcription

## Abstract

Bacterial pathogens use protein secretion systems to transport virulence factors and regulate gene expression. Among pathogenic mycobacteria, including *Mycobacterium tuberculosis* and *Mycobacterium marinum,* ESX-1 (ESAT-6 system 1) secretion is crucial for host interaction. Secretion of protein substrates by the ESX-1 secretion system disrupts phagosomes, allowing mycobacteria cytoplasmic access during macrophage infections. Deletion or mutation of the ESX-1 system attenuates mycobacterial pathogens. Pathogenic mycobacteria respond to the presence or absence of the ESX-1 system in the cytoplasmic membrane by altering transcription. Under laboratory conditions, the EspM repressor and WhiB6 activator control transcription of specific ESX-1-responsive genes, including the ESX-1 substrate genes. However, deleting the *espM* or *whiB6* genes does not phenocopy the deletion of the ESX-1 substrate genes during macrophage infection by *M. marinum*. In this study, we identified EspN, a critical transcription factor whose activity is masked by the EspM repressor under laboratory conditions. In the absence of EspM, EspN activates transcription of *whiB6* and ESX-1 genes both during laboratory growth and during macrophage infection. EspN is also independently required for *M. marinum* growth within and cytolysis of macrophages, similar to the ESX-1 genes, and for disease burden in a zebrafish larval model of infection. These findings suggest that EspN and EspM coordinate to counterbalance the regulation of the ESX-1 system and support mycobacterial pathogenesis.

**Importance:** Pathogenic mycobacteria, which are responsible for tuberculosis and other long-term diseases, use the ESX-1 system to transport proteins that control the host response to infection and promote bacterial survival. In this study, we identify an undescribed transcription factor that controls the expression of ESX-1 genes and is required for both macrophage and animal infection. However, this transcription factor is not the primary regulator of ESX-1 genes under standard laboratory conditions. These findings identify a critical transcription factor that likely controls expression of a major virulence pathway during infection, but whose effect is not detectable with standard laboratory strains and growth conditions.

## Introduction

Diverse pathogens rely on secretion systems to transport virulence factor substrates that promote bacterial survival in different host environments (1, 2). *Mycobacterium tuberculosis*, the causative agent of human tuberculosis, and *Mycobacterium marinum*, a poikilothermic fish pathogen, share a conserved ESAT-6 secretion system (ESX-1) that is required for pathogenesis (3–5). The ESX-1 system is composed of <u>E</u>SX <u>c</u>ore <u>c</u>omponents (Ecc’s) that reside within the mycobacterial cell membrane (CM) and cytoplasm (4, 6–8). During macrophage infection, the ESX-1 system transports protein substrates that are secreted components of the ESX-1 machinery (9) and effectors that promote phagosomal damage (10–13). Phagosomal damage enables the essential interaction between pathogenic mycobacteria and the macrophage cytoplasm, which promotes macrophage death and cell-to-cell spread (12, 14–17).

In several Gram-negative pathogens, transcription is responsive to protein secretion systems (2, 18, 19). We and others have shown that the loss of the membrane associated ESX-1 secretion system in *M. marinum* and *M. tuberculosis* resulted in widespread transcriptional changes (20–22). ESX-1 genes are a subset of the ESX-1 responsive genes. The gene most responsive to ESX-1 was *whiB6*. WhiB6 activates the transcription of ESX-1 substrate genes (20, 23, 24). EspM directly represses *whiB6* gene transcription in the absence of the ESX-1 components in the CM (20, 22). Reduced WhiB6 levels led to reduced ESX-1 substrate gene transcription, preventing accumulation of ESX-1 substrates in the absence of the transport machinery (20). Genetic deletion of the *espM* gene, which is divergently encoded from the *whiB6* gene, derepressed *whiB6* transcription, and increased levels of ESX-1 substrates in *M. marinum* during standard laboratory growth conditions (22).

The deletion of ESX-1 substrate genes attenuates pathogenic mycobacteria in infection models (4, 25–28). Despite regulating transcription of ESX-1 substrate genes, the deletion of *whiB6* or *espM* does not phenocopy the loss of ESX-1 substrate genes in macrophage models of infection (20, 22). Therefore, ESX-1 may be differentially regulated in the laboratory and during infection.

We previously used *lacZ*+ transcriptional fusions to measure EspM and WhiB6-dependent transcription from the *M. marinum whiB6* promoter. In the absence of both EspM and WhiB6, *β*-galactosidase activity was ∼2-fold higher than in the absence of EspM alone (22), suggesting that *whiB6* transcription was activated in the absence of both EspM and WhiB6. We exploited our knowledge of the EspM repressor as a tool to identify an additional regulator of ESX-1 transcription.

## Results

### EspN activates *whiB6* transcription in the absence of EspM

To identify regulators activating *whiB6* transcription in the absence of EspM, we previously used the *whiB6-espM* intergenic region to enrich proteins from lysate from the *ΔespM M. marinum* strain (22). Mass spectrometry identified MMAR_1626 as the only protein enriched for binding the *whiB6-espM* intergenic region in the absence of *espM* [Fig. 1A, original data in SI Table D of (22), analyzed again in SI Dataset SIA-C].

**Figure 1.**
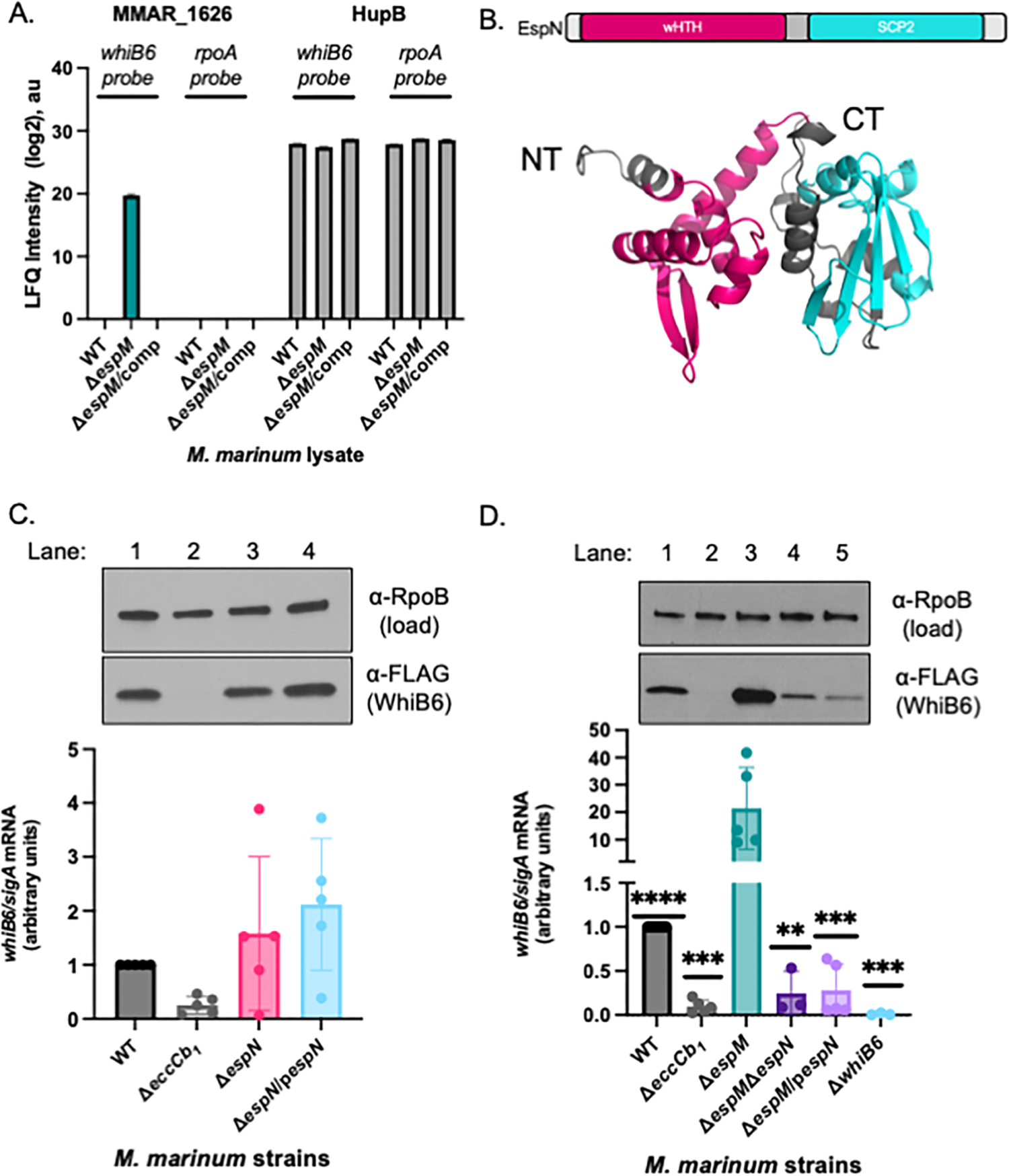
EspN binds the *whiB6* promoter and activates *whiB6* expression in the absence of EspM. **A.** Mass spectrometry analysis of the DNA affinity chromatography showing enrichment of the MMAR_1626 and HupB proteins. The HupB protein binds nonspecifically to both DNA probes. The scale represents the log2 intensity of MS peak arears. a.u. arbitrary units. The data was published in Sanchez et al (22) and was adapted in Dataset S1. **B.** The predicted domain structure of MMAR_1626, which we renamed EspN. Modeled using RoseTTAFold from Robetta (86). Model confidence: 0.80. wHTH: winged helix turn helix, SCP2: sterol carrier protein 2. **C.** (top) Western blot analysis of *M. marinum* cell-associated proteins. RpoB is a loading control. All strains include a *whiB6-3xFl* allele at the *whiB6* locus (20). (bottom) Relative qRT-PCR analysis of *M. marinum* strains compared to *sigA* transcript levels. Statistical analysis was performed using a one-way ANOVA (* *P*=.0359) followed by a Dunnett’s multiple comparisons test relative to the WT strain. **D.** (top) Western blot analysis of 10μg of *M. marinum* whole cell lysates. RpoB serves as a loading control. All strains include a *whiB6-3xFl* allele at the *whiB6* locus (20), (bottom) qRT-PCR of the *whiB6* transcript relative to *sigA*. Significance was determined using a one-way ordinary ANOVA (*P*=.0001), followed by a Tukey’s multiple comparison test. Significance shown is relative to the Δ*espM* strain, with additional statistics of interest discussed in the text. **** *P<.0001, *** P=.*0002 for Δ*eccCb1, ** P*=.0010*, *** P*=.0002 for Δ*espM*/p*espN*, *** *P*=.0009 for Δ*whiB6.* Western blots are representative of three independent biological replicates. All qRT-PCR includes at least three independent biological replicates, each in technical triplicate.

MMAR_1626 is an unstudied putative transcriptional regulator with a predicted N-terminal winged helix-turn-helix domain, and a C-terminal sterol carrier protein 2 (SCP2) domain (Fig. 1B, Fig. S1D) (29, 30). Orthologs are found across *Mycobacterium* and other high G+C Gram-positive bacteria, including members of the TB complex, *M. tuberculosis* (*Rv1725c*) and *M. bovis* (*Mb1754c*), and non-pathogens, including *M. smegmatis* (31, 32). Based on the following data, we propose naming this protein EspN, according to accepted nomenclature (33).

We hypothesized that EspN activates transcription of the *whiB6* gene. To test this hypothesis, we generated an unmarked *espN* deletion strain using allelic exchange (Fig. S1A and S1B) (34). The parental *M. marinum* M strain (wild type, WT) has a *whiB6* allele with a C-terminal 3xFLAG epitope integrated at the *whiB6* locus (20). We generated the complementation strain by integrating a copy of the *espN* gene behind the constitutive mycobacterial optimal promoter at the *attL* site (Fig. S1C). We measured *espN* transcription using qRT-PCR (Fig. S2A). The *espN* transcript was present in the WT strain, but absent from the Δ*espN* strain (*P*=.0002, compared to the WT strain, inset). *espN* transcription was significantly increased (≥40-fold) in the Δ*espN/pespN* strain compared to the WT strain (*P*<.0114, Fig S2A).

To evaluate the role of EspN on *whiB6* transcription, we measured *whiB6* transcript and protein levels in the Δ*espN* and Δ*espN/pespN* strains relative to the WT strain and Δ*eccCb_1_* strains using qRT-PCR (Fig. 1C). We detected *whiB6* transcript and protein in the WT strain (Fig 1C, lane 1). The Δ*eccCb_1_*strain does not produce the EccCb_1_ protein, a cytoplasmic component of the ESX-1 secretion system (4). The loss of EccCb_1_ destabilizes the ESX-1 membrane complex (7, 20, 35), resulting in transcriptional repression of *whiB6* (20, 22). Consequently, the WhiB6-Fl protein was not detected in the Δ*eccCb_1_* strain (Fig. 1C, lane 2) (20). In contrast, *whiB6* transcription in the Δ*espN* and Δ*espN*/*pespN* strains was stochastic, with the mean from three experiments not significantly different from the WT strain. The levels of WhiB6-Fl protein in the Δ*espN* strain and the Δ*espN*/p*espN* strains were similar to the WT strain (Fig. 1C, lanes 3 and 4). We also test the levels of *espF* and *esxA* transcription in the Δ*espN* strain and the Δ*espN*/p*espN* strains. EspF and EsxA are ESX-1 substrates, that are both transcriptionally regulated by WhiB6 (20, 24). Similar to *whiB6* transcript, the *espF* (Fig. S2B) and *esxA* (Fig. S2C) was stochastic. From these data, we conclude that *whiB6* transcription, and the transcription of *espF* and *esxA* was not dependent on EspN under the conditions tested.

We identified both EspM and EspN using DNA affinity chromatography with the *whiB6* promoter DNA (22). EspM was specifically enriched from the WT *M. marinum* lysate, while EspN was only specifically enriched from the *ΔespM* lysate [Fig. 1A, Table D in (22), and Dataset SI, SIA-C). This led us to hypothesize that EspN regulated *whiB6* transcription in the absence of and in opposition to the EspM repressor. To test this, we deleted *espN* from the Δ*espM M. marinum* strain. We also introduced an integrating plasmid constitutively expressing *espN* into the Δ*espM* strain. Using qRT-PCR (Fig. S2D), we found that the *espN* transcript was absent from the Δ*espM*Δ*espN* strain, and significantly increased in the Δ*espM/pespN* strain (∼60-80-fold, *P*<.0001). We measured changes in *whiB6* transcript and protein levels in the Δ*espM*Δ*espN* and Δ*espM*/p*espN* strains compared to the Δ*espM* strain (Fig. 1D). Consistent with our prior work, *whiB6* transcript was significantly reduced in the Δ*eccCb_1_* strain (*P*<.0001) and significantly increased in the Δ*espM* strain relative to the WT strain (*P*<.0001) (20, 22). The WhiB6-Fl protein reflected the differences in transcript levels (lanes 1-3, Fig. 1D) (22). Deletion and overexpression of the *espN* gene in the Δ*espM* strain significantly reduced *whiB6* transcript and WhiB6 protein levels (lanes 4 and 5, Fig. 1D.) compared to the Δ*espM* (*P*=.0002 and *P*=.0009) and WT (*P*<.0001) strains. We conclude that *espN* overexpression in the Δ*espM* strain causes a loss of EspN function, similar to deletion of *espN* (36). Our data support that EspN is required for the elevated *whiB6* transcription in the Δ*espM* strain. Together, our findings suggest that EspN activates *whiB6* transcription in the absence of EspM, working in opposition to EspM at the *whiB6* promoter.

### EspN activates the transcription of ESX-1 component and substrate genes

WhiB6 positively regulates the transcription of several ESX-1 substrate genes (20, 24). We therefore tested how regulation by EspM and EspN impacts ESX-1 function during laboratory growth. *M. marinum* exhibits contact-dependent, ESX-1-dependent hemolytic activity (5, 37). We measured sheep red blood cell (sRBC) lysis to define how EspM and EspN impact ESX-1 activity (Fig. 2A). Water and PBS were “no bacteria” controls, causing total and baseline sRBC lysis as measured by OD_405_, respectively (Fig. 2A). *M. marinum* lysed sRBCs in an ESX-1-dependent manner (Δ*eccCb_1_* vs WT, *P*<.0001). Deletion or overexpression of *espN* did not significantly impact hemolytic activity. The Δ*espM* strain lysed sRBCs slightly less than the WT strain (*P*=.0122). Although the loss of EspM or EspN alone did not greatly impact hemolytic activity, deletion or over-expression of *espN* in the Δ*espM* strain abolished hemolysis (*P*<.0001), similar to the Δ*eccCb_1_* strain. These findings suggest that EspM and EspN are collectively required for ESX-1 dependent lytic activity. Moreover, EspN is essential for lytic activity only in the absence of EspM.

**Figure 2.**
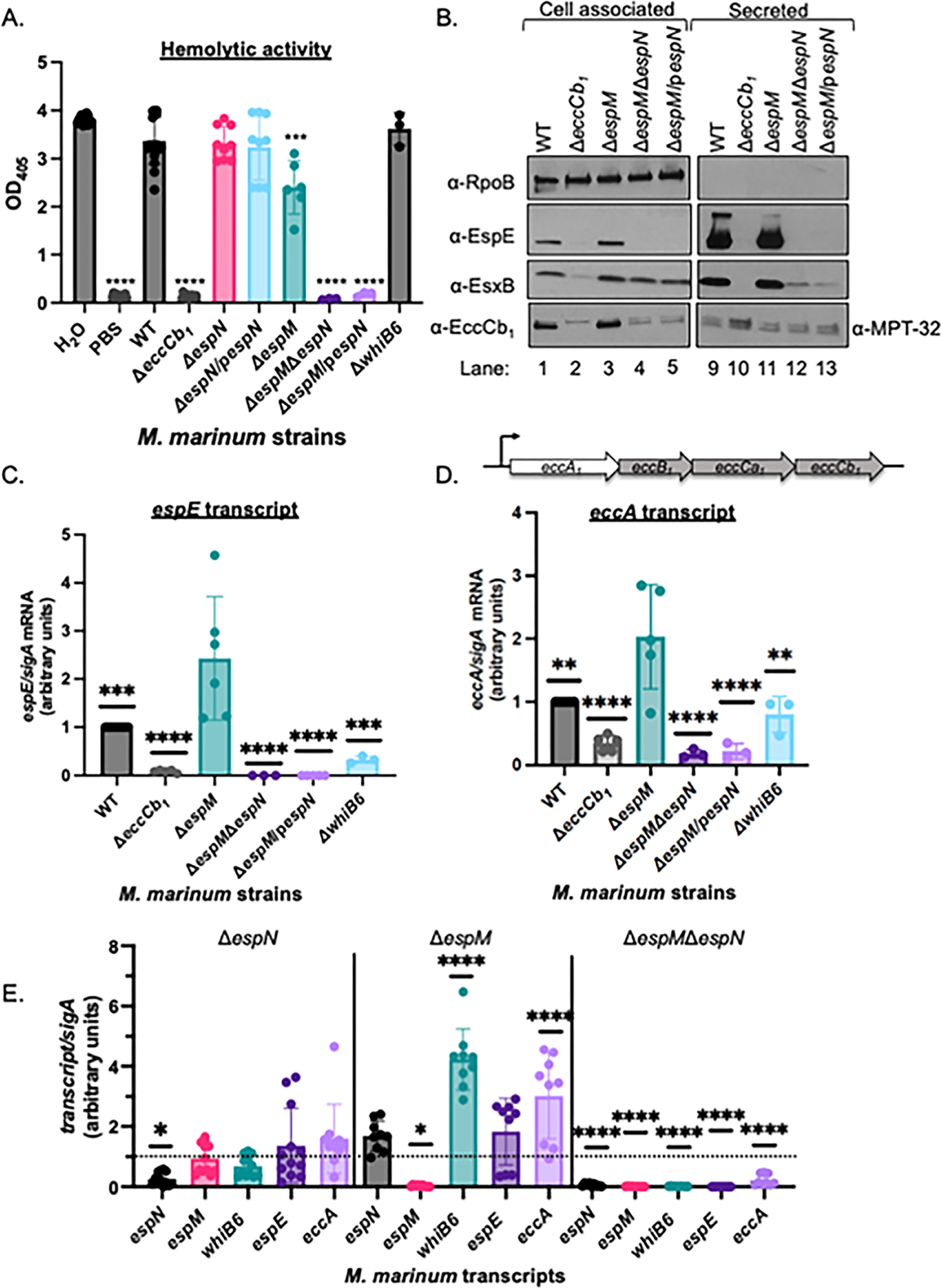
EspN and EspM control transcription of ESX-1 components and substrates. **A.** Sheep red blood cell lysis (sRBC) measuring hemolytic activity of *M. marinum.* Statistical analysis was performed using a one-way ordinary ANOVA followed by a Dunnett’s multiple comparisons test relative to the WT strain. **** *P*<0.0001 *, *** P*=.0003. Data includes three biological replicates each in technical triplicate. **B.** Western blot of 10μg of *M. marinum* cell-associated proteins. All strains include a *whiB6-Fl* allele. RpoB is a control for lysis. MPT-32 is a loading control for the secreted fractions. Blot is representative of three independent biological replicates. **C.** Relative qRT analysis the *espE* transcript compared to *sigA* transcript levels in *M. marinum*. Statistical analysis was performed using a one-way ordinary ANOVA (*P*<.0001) followed by a Tukey’s multiple comparison test. Significance is shown relative to the Δ*espM* strain. *** *P=*0.0010 (WT)*, **** P*<.0001*, *** P=*0.0003 (Δ*whiB6*). **D.** Relative qRT analysis of the *eccA* transcript compared to *sigA* transcript levels in *M. marinum*. Statistical analysis was performed using a one-way ordinary ANOVA (*P*<.0001) followed by a Tukey’s multiple comparison test. Significance shown relative to the Δ*espM* strain. ** *P=*.0012 (WT), *P*=.0026 (Δ*whiB6*)*, **** P*<.0001*, *** P=*0.0003. **E.** Relative qRT-PCR of the *espN, espM, whiB6, espE* and *eccA* transcripts during macrophage infection. RAW 264.7 cells were infected with an MOI of 20, and *M. marinum* strains were isolated at 4 hours post infection. Outliers were identified using ROUT analysis, Q=.05%. Statistical analysis was performed using a one-way ordinary ANOVA (*P=*.0004 for *ΔespN, P*<.0001 for Δ*espM* and Δ*espM*Δ*espN*) followed by a Dunnett’s multiple comparison test relative to the WT strain (dotted line) in each strain. For Δ*espN*, * *P=*.0352, Δ*espM*, * *P*= .0320, **** *P*<.0001; for Δ*espM*Δ*espN*, **** *P*<.0001. For all qRT-PCR, data includes three independent biological replicates each in technical triplicate.

We sought to determine why EspM and EspN were essential for ESX-1 hemolytic activity. Considering that WhiB6 regulates ESX-1 transcription (20, 23, 24), if EspM and EspN were essential for lytic activity solely because they regulate *whiB6* transcription, then we would expect that the hemolytic activity of the Δ*espM*Δ*espN* strain would phenocopy the Δ*whiB6* strain. Consistent with our previous findings, the Δ*whiB6* strain retained hemolytic activity [Fig. 2A, and (20)]. These data indicate that the loss of WhiB6 alone in the Δ*espM*Δ*espN* strain is insufficient to explain the loss of hemolytic activity.

To determine what caused the loss of hemolytic activity of the Δ*espM*Δ*espN* and Δ*espM*/p*espN* strains, we measured the production and secretion of two ESX-1 substrates, EsxB and EspE, using western blot analysis. EsxB is an early substrate and likely secreted component of the ESX-1 system (4, 9). The loss of EsxB abolishes ESX-1 substrate secretion and hemolytic activity (5, 28). EspE, a late substrate abolishes hemolytic activity, but does not affect ESX-1 substrate secretion, except EspF (9, 28). EsxB and EspE were produced in (Fig. 2B, lane 1, αEsxB, αEspE) and secreted from (lane 9) the WT strain during *in vitro* growth. The Δ*eccCb_1_* strain had reduced EsxB and EspE levels (lane 2), due to EspM-dependent repression of *whiB6* transcription (22). Neither protein was secreted (lane 10) due to a loss of the ESX-1 membrane components. EspE and EsxB were produced (lane 3) and secreted from the Δ*espM* strain (lane 11), consistent with our prior findings (22). Deletion or overexpression of the *espN* gene in the Δ*espM* strain abolished EspE production (lanes 4 and 5) and therefore secretion (lanes 12 and 13). EsxB was produced in both strains, but EsxB secretion was reduced compared to the WT and Δ*espM* strains (lanes 12 and 13).

Reduced EsxB secretion could be explained by reduced ESX-1 components in the CM. EccCb_1_ is required for the stability of the ESX-1 complex (20). As measured by western blot analysis (Fig. 2B), EccCb_1_ protein was detected in the cell-associated proteins from the WT strain and absent from the Δ*eccCb_1_* strain (lanes 1 and 2, lower band). The Δ*espM* strain produced EccCb_1_ protein (lane 3,). In the absence of EspM and EspN, EccCb_1_ protein levels were lower than the WT strain (lanes 4 and 5).

The loss of EspE protein and reduced EccCb_1_ protein could be explained by reduced transcription. We tested if EspN regulated the transcription of *espE* or the ESX-1 component genes when EspM was absent. As measured by qRT-PCR (Fig. 2C) *eccCb_1_* deletion significantly reduced (*P*<.0001) and *espM* deletion significantly increased (*P*=.0010) *espE* transcription relative to the WT strain, consistent with Fig. 2B and our previous findings (22, 38). Deletion or overexpression of *espN* in the Δ*espM* strain abolished *espE* transcription, significantly different from the Δ*espM* (*P*<.0001) and the WT strains (*P*<.0001). Importantly, *whiB6* deletion significantly reduced (*P=*.0003), but did not abolish *espE* transcript. Therefore, the loss of WhiB6 was insufficient to cause the loss of *espE* transcription in the Δ*espM*Δ*espN* strain. We conclude that EspN is required for transcription of *espE* in the Δ*espM* strain (Fig. 2C).

The *eccCb_1_* gene is downstream of three other ESX-1 component genes (Fig. 2D, *eccA, eccB, eccCa_1_).* It is not known if the *ecc* genes are operonic. To test if reduced EccCb_1_ protein was due to reduced *ecc* transcription, we measured *eccA* transcription using qRT-PCR (Fig. 2D). *eccA* transcript was significantly reduced (*P*<.0001) in the Δ*eccCb_1_* strain, and significantly increased (*P*=.0012) in the Δ*espM* strain as compared to the WT strain. Together, these data suggest that EspM represses *eccA* transcription. Deleting or overexpressing *espN* in the *ΔespM* strain significantly reduced *eccA* transcription compared to the Δ*espM* (*P*<.0001) and WT strains (*P*<.0001). Interestingly, while *eccA* transcript in the Δ*whiB6* strain was significantly reduced compared to the Δ*espM* strain (*P=.0026*), it was not significantly different from the WT strain. These findings suggest that EspN activates and EspM represses *eccA* transcription independently of WhiB6.

To test how EspN and EspM impact transcription during infection, we infected RAW 264.7 macrophages with *M. marinum* strains lacking *espM, espN* or both *espM* and *espN. M. marinum* can escape the phagosome between 2 and 4 hours post infection [hpi, (39)]. Since ESX-1 functions in the phagosome, we harvested the bacteria at 4 hpi and used qRT-PCR to measure *whiB6, espN, espM, espE* and *eccA* transcripts in *M. marinum. whiB6, espM, espE* and *eccA* transcription was not significantly different between the WT and Δ*espN* strains (Fig. 2E), in agreement with Fig. 1C. Similar to our findings during laboratory growth, deletion of *espM* significantly increased *whiB6* (*P*<.0001) and *eccA* (*P*<.0001) transcript levels relative to the WT strain. The *espN* and *espE* transcripts were also higher than the WT strain, but did not reach significance. Strikingly, all five transcripts were significantly reduced in the Δ*espM*Δ*espN* strain compared to the WT strain. These data suggest that EspM and EspN jointly regulate *whiB6, espE,* and *eccA* transcription during macrophage infection under the conditions tested.

### EspN is required during *M. marinum* infection of macrophages and zebrafish

Our data propose a model where EspM and EspN counterbalance to regulate ESX-1 gene transcription, in both the lab and during infection. However, EspM obscures the regulatory role of EspN under laboratory conditions and during macrophage infection. *M. marinum* cause macrophage cytolysis in an ESX-1dependent manner. Strains lacking the ESX-1 system remain in the phagosome and are non-cytolytic (10). We infected RAW264.7 cells with *M. marinum* (MOI of 4) and imaged 24 hpi using ethidium homodimer (EthD-1) staining to measure macrophage cytotoxicity (Fig. 3A). EthD-1 selectively stains DNA in cells with permeabilized cell membranes, reflecting the cytolytic activity of *M. marinum* (5, 40). Infection with WT *M. marinum* led to macrophage cytotoxicity (Fig. 3A). Uninfected and Δ*eccCb_1_* infected macrophages exhibited significantly less cytotoxicity than WT-infected macrophages (*P*<.0001). Infection with the Δ*espN* strain significantly decreased macrophage cytotoxicity, similar to the Δ*eccCb_1_* strain (*P*<.0001). Overexpressing *espN* in the Δ*espN* strain (*P*<.0001) partially complemented cytolytic activity. The Δ*espMΔespN* and Δ*espM*/p*espN* strains were non-cytolytic, similar to the *ΔespN* and *ΔeccCb_1_* strains. From these data we conclude that EspN is independently essential for *M. marinum* to cause macrophage cytotoxicity.

**Figure 3.**
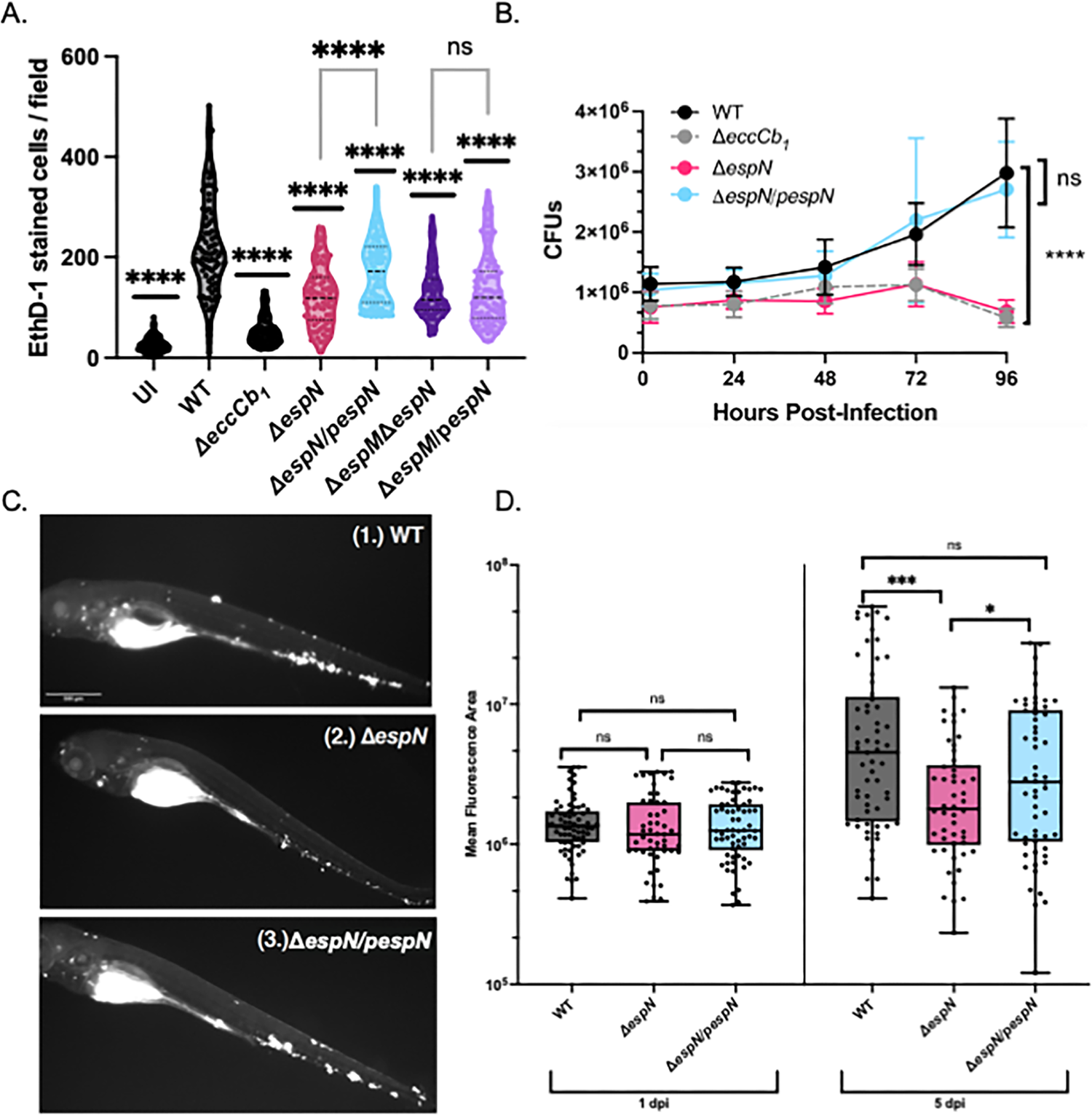
EspN is required for pathogenesis. **A.** Colony forming units (CFU) of *M. marinum* strains MOI = 0.2. The data points are the average of three independent biological replicates. Significance was determined using an ordinary 2 way ANOVA (*P*<.0001) followed by a Tukey’s multiple comparison test compared to the WT strain. The significance shown is compared to the WT strain at 96-hours post infection (*P*<.0001). The Δ*eccCb_1_* and Δ*espN* strains were also significantly different from the WT strain at 72 hpi (*P*=.0021 and *P*=.0024, respectively). **B.** *M. marinum* burden in zebrafish infection measured using bacterial mCerulean fluorescence. Data are comprised of two biological replicates with 20-30 independent infections per replicate. Statistical analyses were performed using one-way ANOVA followed by a Tukey multiple comparison of each group to the WT strain (*** *P*=.00012; * *P*=.021). **C.** Representative images of zebrafish infected with an initial dose of 150-200 fluorescent bacilli for (1.) WT, (2.) Δ*espN*, or (3.) Δ*espN*/p*espN* at 5 days post-infection. Scale bar is 500μm. **D.** Macrophage cytolysis as measured by EthD-1 staining 24h post-infection with *M. marinum* at an MOI of 4. Statistical analysis was performed using a one-way ANOVA followed by a Dunnett’s multiple comparisons test relative to the WT strain. (**** *P*<.0001) Each dot represents the number of EthD-1-stained cells in a single field. A total of 10 fields were counted using ImageJ for each well. Processing of 3 wells was performed for each biological replicate. A total of 90 fields were counted for each strain.

To test if *espN* was required for mycobacterial growth in macrophages, we performed CFU analysis (Fig. 3B). WT *M. marinum* grew within macrophages during infection. In contrast, the Δ*eccCb_1_* strain was attenuated for growth in macrophages, resulting in CFUs significantly different from the WT strain at 72 and 96 hpi. Consistent with the cytotoxicity data, the Δ*espN* strain was attenuated for growth in the macrophages, similar to the Δ*eccCb_1_* strain. Overexpression of the *espN* gene restored growth of *M. marinum* similar to the WT strain. From these data we conclude that EspN is required for *M. marinum* growth and during macrophage infection.

To assess EspN’s importance during animal infection, we made constitutively fluorescent versions of the WT, *ΔespN,* and complemented *M. marinum* strains. Using the zebrafish larval model of mycobacterial infection, we monitored burden over five days of infection as measured by a validated Fluorescence Pixel Count (FPC) assay [Fig. 3C and 3D, (41, 42)]. Bacterial burden was substantially reduced at five days post-infection. We found that *in vivo*, *ΔespN* had 3.4-fold reduced burden compared to WT at five days post-infection (*P*=.00012) and was complemented by constitutive expression of the *espN* gene (Fig. 3C and 3D). From these data, we conclude that EspN is required during zebrafish infection by *M. marinum.* Together our data support that EspN is essential for mycobacterial infection.

### Overexpression of EspN causes loss of ESX-1 function

We found it curious that *espN* overexpression in the Δ*espM* strain phenocopied the Δ*espM*Δ*espN* strain during laboratory growth and macrophage infection. Our data argue against an unidentified mutation as the reason for the shared phenotypes. *espM* expression in the Δ*espM* strain complements all phenotypes associated with EspM loss. We performed whole genome DNA sequencing on the WT, Δ*espM*, Δ*espN*, Δ*espM*Δ*espN* and the Δ*espM*/p*espN* strains (Dataset S2). No consistent mutations were identified between the Δ*espM*Δ*espN* and Δ*espM*/p*espN* strains that could account for the observed phenotypes.

We hypothesized that *espN* overexpression was affecting endogenous EspN activity. We designed mutations in the mycobacterial optimal promoter (mop) focusing on −7 and −12 positions (Fig. 4A). We confirmed the mutations by DNA sequencing, introduced the p*espN* plasmids into the Δ*espM* or *ΔespMΔespN M. marinum* strains and measured *espN* expression using qRT-PCR. The mutated plasmids resulted in significantly reduced *espN* expression compared to the parental plasmid (Fig. 4B, WT, *P*<.0001). Although overexpression of *espN* abolished the hemolytic activity of the Δ*espM* strain (*P*<.0001), reduced *espN* expression did not significantly alter hemolysis of the Δ*espM* strain (Fig. 4C). From these data, we conclude that the *espN* overexpression caused the loss of hemolysis in the Δ*espM* strain. Notably, reduced *espN* expression from the mutated promoter did not restore hemolysis of the Δ*espM*Δ*espN* strain (Fig. 4C).

**Figure 4.**
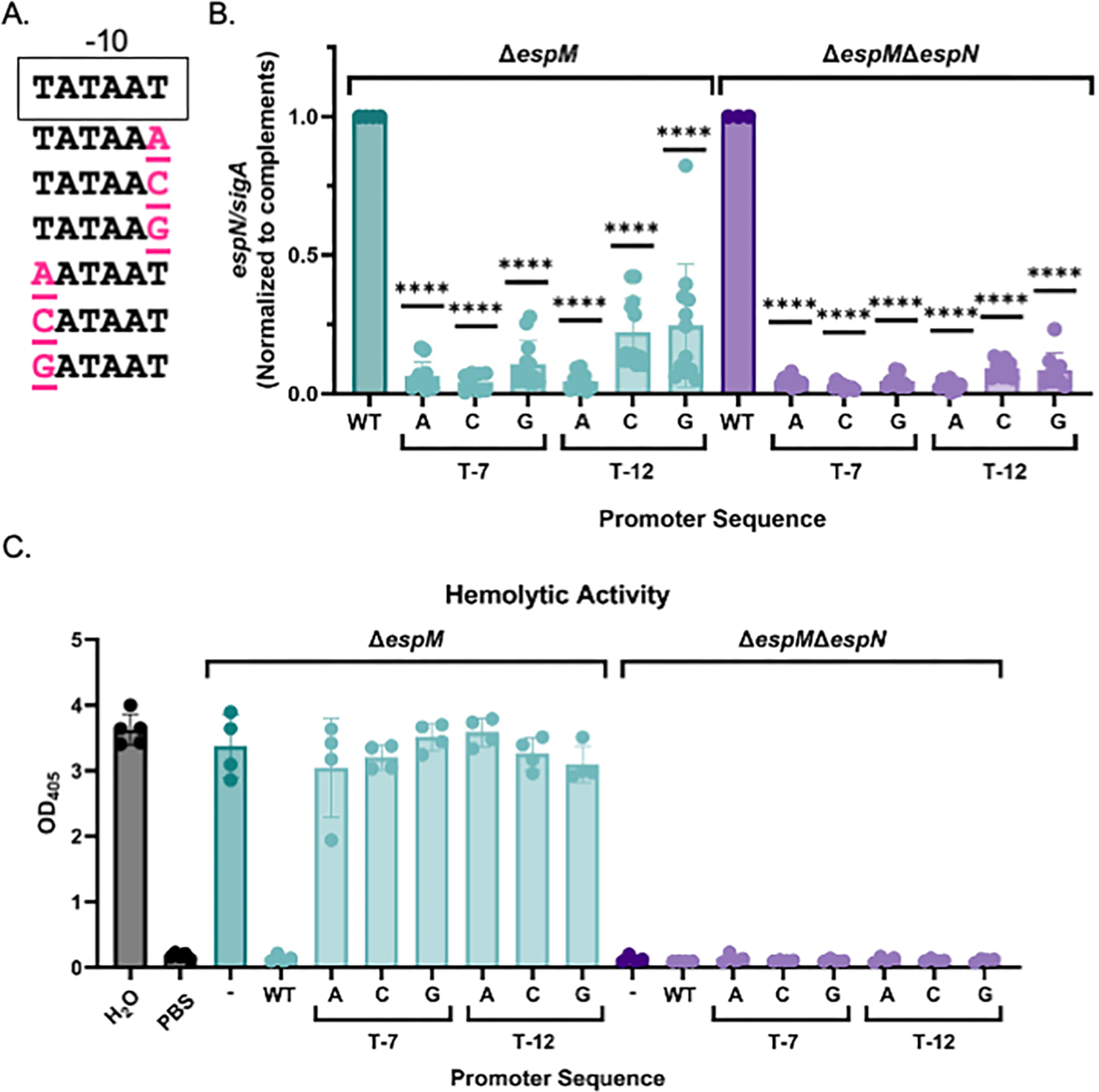
High levels of EspN transcription are required for dominance in the Δ*espM* strain. **A.** Schematic of the −10 region from the mycobacterial optimal promoter driving *espN* transcription. Pink residues are mutations in the −7 and −12 positions. **B.** Relative qRT analysis the *espN* transcript compared to *sigA* transcript levels in *M. marinum*. Outliers were identified using ROUT analysis, Q=0.5%. Statistical analysis was performed using a one-way ordinary ANOVA (*P*<.0001) followed by a Dunnett’s multiple comparison test. Significance is shown relative to the Δ*espM/pespN* or Δ*espM*Δ*espN*/p*espN* strain. *\*\*\*\* P*<.0001. Data includes three biological replicates each in technical triplicate. **C.** sRBC lysis measuring hemolytic activity of *M. marinum.* Statistical analysis was performed using a one-way ordinary ANOVA followed by a Dunnett’s multiple comparisons test relative to the Δ*espM* or Δ*espM*Δ*espN* strain. **** *P*<.0001. Data includes three biological replicates each in technical triplicate.

### EspE and the N-terminus of EspM are linked to EspN function

To understand the transcriptional network (Fig. 5A), we examined regulatory interactions between EspM and EspN. Assessing *espN* transcription in the Δ*espM* strain, and *espM* transcription in the Δ*espN* strain (Fig. S4A and B) revealed no significant changes. We conclude that EspM and EspN do not regulate each other transcriptionally under the conditions tested.

**Figure 5.**
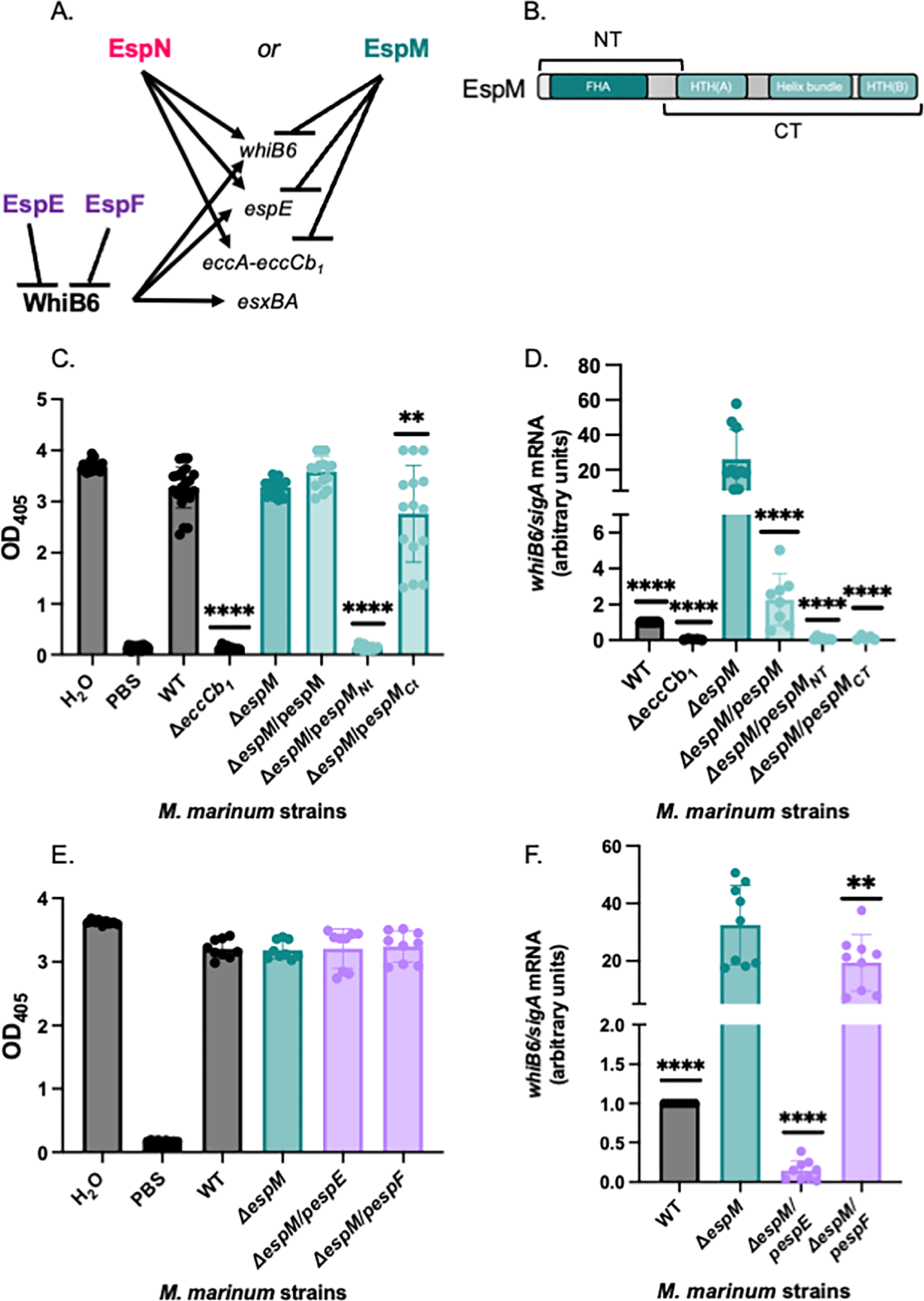
Overexpression of the EspM N-terminus or EspE negatively impacts ESX-1 transcription in the absence of EspM. **A.** Schematic of transcriptional regulation by EspM, EspN and WhiB6. EspE and EspF are ESX-1 substrates that negatively regulate the WhiB6 transcription factor (28). **B.** Schematic of the predicted domains of the EspM protein. NT: N-terminus, FHA: Forkhead associated domain, HTH: Helix turn helix. **C.** sRBC lysis measuring hemolytic activity of *M. marinum.* Outliers were identified using ROUT analysis, Q=.05%. Statistical analysis was performed using a one-way ANOVA (*P*<.0001) followed by a Dunnett’s multiple comparisons test (**, *P*=.0034, **** *P<*.0001). The data includes at least three independent biological replicates each in technical triplicate. **D.** Relative qRT-PCR analysis of *whiB6* compared to *sigA* transcript levels. Significance was determined using a one-way ordinary ANOVA (P<.0001), followed by a Bennett’s multiple comparison test (**** *P<*.0001) relative to the Δ*espM* strain. The qRT-PCR data includes at least three independent biological replicates each in technical triplicate. **E.** sRBC lysis measuring hemolytic activity of *M. marinum.* Outliers were identified using ROUT analysis, Q=.05%. Statistical analysis was performed using a one-way ordinary ANOVA (*P*=.9639), which did not indicate significant differences. The data includes at least three independent biological replicates each in technical triplicate. **F.** Relative qRT-PCR analysis of *whiB6* compared to *sigA* transcript levels. Outliers were identified using a ROUT analysis (Q=0.5%). Significance was determined using a one-way ordinary ANOVA (P<.0001), followed by a Bennett’s multiple comparison test (**, *P*=.0070, **** *P<*.0001) relative to the Δ*espM* strain. The qRT-PCR data includes at least three independent biological replicates each in technical triplicate.

Because EspN overexpression abolished hemolytic activity and ESX-1 gene expression in the Δ*espM* strain, we asked if overexpressing other regulators impacted ESX-1 function in the absence of EspM. EspM has an N-terminal forkhead-associated (FHA) domain (EspM_NT_, Fig. 5B) and two helix-turn-helix (HTH) domains between a helical bundle (EspM_CT_ in Fig. 5B) at the C-terminus. We constitutively expressed the *espM, espM_NT_* and *espM_CT_*genes including a C-terminal V5 epitope tag in the Δ*espM* strain. All EspM-V5 proteins were expressed in *M. marinum* (lanes 4-6, Fig. S5A)*. espM-V5* expression in the Δ*espM* strain did not significantly impact hemolytic activity (Fig. 5C). *espM_NT_-V5* expression abrogated hemolytic activity of the Δ*espM* strain (*P<.0001*). *espM_CT_-V5* expression in the Δ*espM* strain significantly impacted hemolytic activity (*P=.0034*), causing stochasticity relative to the other strains. These data suggested that the Δ*espM/pespM_NT_*-V5 strain effectively phenocopied the Δ*espM*Δ*espN* strain regarding ESX-1-dependent hemolytic activity (Fig. 3B).

Overexpressing the EspM-V5 or the EspM_CT_-V5 protein significantly reduced the *whiB6* (*P*=.0014, *P*=.0050, Fig. 5C)*, eccA* and *espE* transcripts (Fig. S5B-D) relative to the Δ*espM* strain. Overexpression of EspM-V5 resulted in reduced WhiB6Fl, EccCb_1_, EspE, and EsxB protein levels compared to the Δ*espM* strain (Fig. S5A). Expression of the *espM_CT_* gene resulted in a loss of detectable WhiB6-Fl protein and further reductions of the EccCb_1_, EspE and EsxB proteins compared to the Δ*espM* and Δ*espM/pespMV5* strains (Fig. S5A). The EspM_CT_ binds the *whiB6* promoter and represses *whiB6* transcription in *M. marinum* (22, 43). Although EspM_NT_ does not bind the *whiB6* promoter (22), expression of *espM_NT_* in the Δ*espM* strain significantly reduced *whiB6* (*P*=.0015), *espE* (*P*<.0001), and *eccA* (*P*<.0001) transcription, resulting in a loss of EspE and reduced WhiB6-Fl and EccCb_1_ proteins compared to the Δ*espM* and complemented strains (Fig. 5C and Fig. S5A, statistical analyses in Fig S5). None of the strains tested exhibited significant reductions in *espN* transcription (Fig. S5E).

EspE and EspF are dual functioning ESX-1 substrates. Both proteins negatively regulate WhiB6 activity in *M. marinum*, and their secretion is required for hemolytic activity and for virulence (28). Overexpressing *espE* or *espF* did not impact hemolytic activity (Fig. 4E) but significantly reduced *whiB6* transcription in the Δ*espM* strain. However, while *espE* overexpression reduced *whiB6* transcription to levels below the WT strain, overexpression of *espF* resulted in transcription levels ∼20x higher than the WT strain. From these data, we conclude that overexpression of EspE and EspM_NT_, but not EspF, results in phenotypes consistent with EspN loss of function in the absence of EspM.

## Discussion

Our prior studies hinted at the existence of an additional ESX-1 activator (22). Here, we discovered and characterized EspN, a transcriptional regulator of the ESX-1 system that is essential for infection. Our data supports that EspN and EspM function as a switch that regulates the transcription of ESX-1 genes. When the ESX-1 system is absent, EspM represses *whiB6,* ESX-1 component (EccA and others) and substrate genes, preventing substrate accumulation in the absence of secretion (20, 22). EspM also regulates the transcription of additional genes in *M. marinum* (22). In the presence of the ESX-1 system, WhiB6 activates ESX-1 substrate gene transcription and other genes, allowing production of substrates during active secretion (20, 23, 24). The presence of EspM masked a role for EspN in regulating ESX-1 gene transcription. However, in the absence of EspM, EspN is essential for ESX-1 activity likely because it regulates *espE* transcription and contributes to *eccA* transcription, optimizing ESX-1 component production and substrate secretion. Deleting *espN* attenuated the cytolytic activity and growth of *M. marinum* during macrophage infection, similar to a loss of ESX-1. EspN was likewise necessary for robust infection in zebrafish larvae. Our findings support that EspN is required for ESX-1 function during infection possibly due to transcriptional regulation of ESX-1 genes. Deleting both *espM* and *espN* from *M. marinum* abolished cytolytic activity and *whiB6, espE* and *eccA* transcription during macrophage infection, which differed from deleting *espN* alone. Finally, overexpression of *espN, espM_NT_*and *espE* specifically disrupted ESX-1 regulation, demonstrating connections in the transcriptional network. Together, these data demonstrate a critical role for EspN and the ESX-1 transcriptional network during infection.

Our data support that EspN may function independently and together with EspM to regulate transcription and to promote pathogenesis. *espN* deletion did not have regulatory phenotypes under laboratory conditions and during macrophage infection under the conditions we tested. However, the Δ*espN* strain was attenuated both in macrophage and in zebrafish larvae. EspN-dependent regulation of ESX-1 or additional genes may be essential for virulence. The deletion of both EspN and EspM abolished ESX-1 transcription under lab conditions and during macrophage infection, which differed from deleting either regulator alone. Defining the regulatory targets of EspN, and how the EspM and EspN regulons compare, will likely require measuring global gene expression during specific stages of infection.

Some of our strains exhibited variable transcript levels (the Δ*espN* and Δ*espM* strains, Fig. 1C-D, Fig. 2D-E, and Fig. 5C-D) or hemolytic activity (Δ*espM*, Fig. 2A, Δ*espM*/p*espM_CT_*, Fig. 5F) across multiple experiments. Our studies relied on population averages for each assay. In regulatory systems such as Type III secretion-dependent transcriptional regulation in *P. aeruginosa*, persister cells in *E. coli,* and cannibalism and competence in *B. subtilis*, stochasticity reflects that individual cells within a population have different gene expression patterns (44, 45). This “bistability” indicates a switch between two states, rather than intermediate ones (44). The stochasticity of our results may suggest that ESX-1 regulatory network includes a bistable switch that operates early during infection, possibly through pairs of mutually exclusive regulators or positive autoregulation (44). While the molecular nature of the switch is unknown, EspM and EspN likely cannot simultaneously occupy the *whiB6, eccA* or *espE* promoters. Host-specific signals may regulate the switch including oxidative stress, pH and other phagosomal cues. *espN* is divergently transcribed from a putative methyl-transferase gene (*MMAR_1627*) (31). We do not yet know if the genes surrounding *espN* are important for regulation of ESX-1, methylation has been shown to control bacterial genetic switches (46, 47).

ESX-1 expression varies in different strains under laboratory conditions (24). Consistent with our model, ESX-1 gene expression is upregulated in the host (48), and transcriptomic studies in *M. marinum* suggested differential regulation of virulence genes in a variety of environments (49). The differential regulation of protein secretion systems under laboratory conditions and in the host is a common theme in protein secretion in bacterial pathogens. For example, some clinical *Vibrio cholerae* strains have active Type VI systems under laboratory conditions, while the pandemic strains only activate their Type VI systems in the host (50). The specialized Type III secretion systems in *Vibrio* species are regulated by bile salts (51), and by Ca^2+^ and host cell contact in *Pseudomonas* (52–54).

Our findings suggest that EspN activity or expression might respond to host-specific signals. Notably, transcriptional profiling of *M. tuberculosis* from infected macrophages isolated from mouse infections revealed significant upregulation of *espN* (Rv1725c) transcript compared to broth-grown culture (55). At one week post infection in a mouse intravenous model, a TnSeq screen identified a ∼2-fold decrease in representation of transposon insertions in *M. tuberculosis Rv1725c* [*P*=.0039, (56)]. However, at four weeks and eight weeks this effect was no longer present (56, 57). Together, these data suggest that complex regulatory circuits and genetic requirements operate at discrete stages of infection. EspN’s C-terminal SCP2 domain (Fig. 1B) could differentially localize the protein in the mycobacterial cell in response to the host environment (58, 59) to mediate interaction with the mycobacterial cell membrane under specific conditions. SCP-2 domains also mediate lipid transfer of sterols such as cholesterol and fatty acids (60, 61), which are an energy source for mycobacterial pathogens during infection (62, 63). Focusing on EspN’s SCP-2 domain may reveal how EspN senses and responds to the host environment.

Gene dosage is important for biological function across organisms. Increases in copy number or expression of wild-type genes can cause mutant phenotypes, such as aggregation or mislocalization (36). We were surprised that the overexpression of *espN, espM_NT_* and *espE* in the absence of *espM* resulted in the same regulatory phenotypes in *M. marinum* as deleting *espN* from the Δ*espM* strain*. espN, espM_NT_* or *espE* overexpression could result in EspN aggregation or mislocalization. Transcription factor multimerization is a regulatory mechanism in bacterial gene expression (64, 65). In higher organisms, transcription factors can form aggregate-like bodies that serve as functional regulatory mechanisms which mimic loss of function (66). EspN may form multimeric aggregates that prevent DNA interaction. Alternatively, EspN may be mislocalize under overexpression conditions. The EspM_NT_ is a predicted FHA domain. Proteins with FHA domains regulate other Gram-negative secretion systems post-transcriptionally (67–69). Our data also support that EspM may be processed *in vitro*. EspM cleavage might remove it from the promoter, similar to the cI repressor from λ phage (64). Alternatively, cleavage may liberate the N-terminus to regulate EspN activity, allowing the EspM_CT_ to bind DNA and repress gene expression.

Our studies were limited by an inability to complement *espN* expression in the Δ*espM*Δ*espN* strain. We tested overexpression, endogenous and inducible promoters to restore *espN* transcription in the Δ*espM*Δ*espN* strain. We were unable to restore transcription to the levels of the Δ*espM* strain. We suspect that architecture or chromosomal location of *espN* may be important for proper regulation and expression.

Several additional transcription factors regulate *whiB6* and the *espACD* operon which encodes secreted substrates essential for ESX-1 secretion in *M. tuberculosis* (25, 70–72). PhoPR and MprAB regulate *whiB6* expression directly and through the EspR regulator (73–77), in response to pH or stress, both of which are important for survival in the acidified phagosome (78–80). However, the *espACD* operon is dispensable for ESX-1 secretion and pathogenesis in *M. marinum,* and regulation of *whiB6* by PhoPR has not been reported (9, 81). Although this may suggest divergence in ESX-1 regulation between *M. marinum* and in *M. tuberculosis,* the ESX-1 transcriptional network is conserved (20, 21, 82). Widespread ESX-1-dependent gene expression has been reported in both mycobacterial species (20, 21, 82). Moreover, we showed that EspM and the regulatory substrates, EspE and EspF, are functionally conserved in *M. tuberculosis* (22, 28). Future research will be aimed at defining the ESX-1 transcriptional network in *M. tuberculosis*.

Overall, this study further defined the regulatory network underlying control of the ESX-1 secretion system and demonstrated its importance during infection. We have, for the first time, identified an infection-dependent transcriptional activator responsible for regulating both the ESX-1 components and substrates. Our study will serve as a foundation for understanding the molecular complexities of ESX-1 regulation in the host. We are now poised to define the molecular mechanisms underlying how the ESX-1 system senses and responds to a changing host environment.

## Materials and Methods

Bacterial strains were derived from the *M. marinum* M parental strain (ATCC BAA-535), and were maintained as previously described (20, 22, 28, 83). Nomenclature follows the conventions proposed by Bitter et al (33). Genetic deletions were performed using allelic exchange as previously described (20, 22, 28, 83, 84). Hemolytic activity was measured against sRBCs as described previously (20, 22, 28, 83). Cell-associated and secreted mycobacterial proteins were isolated and analyzed as described in (83, 85). Protein levels were measured using western blot analysis. RNA extraction was performed from *M. marinum* using the Qiagen RNeasy kit, followed by qRT-PCR relative to the levels of *sigA* as described previously (22, 43). Macrophage (RAW 264.7) cytotoxicity was measured using Ethidium homodimer uptake following infection by *M. marinum* as described in (38, 83). Mycobacterial CFUs were performed similarly to (85). Protein modeling was performed using Robetta and Pfam as indicated. Zebrafish larvae infections with *M. marinum* were performed and bacterial burden was measured using fluorescent pixel counts as (41). Statistical analysis was performed using GraphPad Prism v. 9 or R within the latest version of R Studio IDE. Detailed methods are in the Supplementary Material.

## Acknowledgments

K.R.N was supported by an Eck Institute for Global Health Fellowship and an Arthur J. Schmitt Fellowship. P.A.C. is supported by the National Institutes of Health under award numbers AI156229, AI106872, AI149147, and AI149235. Support for D.M.T. through NIH award numbers AI130236, AI125517, and AI166304. We thank the Champion Lab for their critical reading of this manuscript. We thank Dr. Robert Abramovitch for his suggestions regarding *M. marinum* RNA isolation during macrophage infection. We thank Dirk Schappinger for his advice on mutations to weaken *espN* overexpression. The content of this article is solely our responsibility and does not necessarily represent the official views of the National Institutes of Health.

## Supplemental Legends

**Fig. S1. Genotypic confirmation of espN deletion and complementation. A.** PCR of upstream and downstream of the *espN* gene using OKN93 and OKN94 primers to confirm genetic deletion in *M. marinum* strains. WT = 1283 bp; Δ*espN* = 665 bp. *M. marinum* genomic DNA (gDNA) serves as a negative control. **B.** Schematic of the *espN* deletion using allelic exchange. **C.** PCR of p*espN* plasmid in *M. marinum* strains using OKN139 and MOPF primers. A 775 bp band indicates the p*espN* is present. *M. marinum* gDNA serves as a negative control. **D.** Schematic of the genetic locus including *espN* in *M. marinum*.

**Fig. S2. EspN and ESX-1 substrate expression in deletion and overexpression *M. marinum* strains.** A. Relative qRT analysis of *espN* in *M. marinum* strains compared to *sigA* transcript levels. Statistical analysis was performed using a one way-ordinary ANOVA *(P*=.0092), followed by a Dunnett’s multiple comparison test. * *P*=.0114. Inset: Zoom in of the comparison between the WT and Δ*espN* strain. The levels of *espN* were compared using an unpaired student’s t-test. *** *P*=.0002. B. *espF* in *M. marinum* strains compared to *sigA* transcript levels. Statistical analysis was performed using a one way-ordinary ANOVA (*P*=.0008), followed by a Dunnett’s multiple comparison’s test. * *P*=0.0264 and C. *esxA* in *M. marinum* strains compared to *sigA* transcript levels. Statistical analysis was performed using a one way-ordinary ANOVA (*P*=.0002), followed by a Dunnett’s multiple comparison’s test. ** *P*=0.0063. D. Relative qRT analysis of *espN* in *M. marinum* strains compared to *sigA* transcript levels. Statistical analysis was performed using a one way-ordinary ANOVA (*P*<.0001), followed by a Dunnett’s multiple comparison’s test. **** *P*<0.0001. Inset: Comparison between the WT and Δ*espM*Δ*espN* strains. The levels of *espN* were compared using an unpaired student’s t-test. **** *P*<0.0001.

**Fig. S3. ESX-1 membrane complex transcripts are regulated by ESX-1. A.** Log_2_ fold-change of ESX-1 membrane complex component transcripts from RNA sequencing data published in Sanchez et al (22). The *eccA-eccCb_1_* transcripts were significantly upregulated in the Δ*espM* strain compared to the overexpression strain (*P = 1.07e-23, 1.11e-19, 3.99e-15, 2.13e-16*). The *eccD_1_* and *eccE_1_* transcripts were not significantly different between the two strains by RNA sequencing. **B.** Relative qRT analysis of *eccA* transcript in *M. marinum* strains compared to *sigA* transcript levels. Statistical analysis was performed using a one way-ordinary ANOVA *(P*=.0001), followed by a Tukey’s multiple comparison test. *** *P*=.0005, **** *P*<.0001.

**Fig. S4: EspM and EspN do not regulate each other transcriptionally in *M. marinum* under laboratory conditions**. qRT PCR on total RNA extracted from *M. marinum* following growth *in vitro* measuring **A.** *espM* transcript or **B.** *espN* transcript relative to *sigA*. None of the measured changes of the *espM* transcript in panel A are significantly different from the WT strain based on a one-way ordinary ANOVA. In panel B, significance was determined using a one-way ordinary ANOVA (*P<.0001*), followed by a Dunnett’s multiple comparison test. **** *P*<.0001 compared to the WT strain. In the inset, the Δ*espM*/*pespN* strain was excluded. In the inset, an unpaired t-test was performed between the WT and the Δ*espM*Δ*espN* and WT strains (*P*<.0001).

**Fig. S5: Expression of EspMNT impacts ESX-1 gene expression.** A) Western blot of *M. marinum* whole cell lysates. All strains include a whiB6-Fl allele. EspM complementation strains are tagged at the C-terminus with a V5 tag. RpoB is a loading control. EspE and EsxB are ESX-1 substrates. B) Relative qRT-PCR analysis of M. *marinum* strains compared to *sigA* transcript levels. The values were normalized to those in the Δ*espM* strain and plotted as the log2 fold change rela-tive to the Δ*espM* strain as a heat map. The qRT-PCR data plotted in this heat map include at least three independent biological replicates each in technical triplicate, are shown in Fig. 5D, and panels C) *eccA*, D) *espE,* and E) *espN* relative to *sigA* in this figure. Each data point is the average of three technical replicates. Statistical analysis was performed using a one-way ordinary ANOVA followed by a Dunnett’s post-hoc test vs the Δ*espM* strain. C) ANOVA *P<*.0001, *** (vs WT, *P*=.0005, vs Δ*espM/pespM, P*=.0003), **** *P* <.0001, D) ANOVA *P<*.0001, \*\**P*=.0068, *** *P*=.0002, **** *P* <.0001, E) ANOVA *P* value was not significant.

**Table S1.** Strains and Plasmids used in this study.

**Table S2.** List of Primers used in this study.

**Dataset S1.** Proteomics Dataset. The dataset is reprocessed data from Ref. (22).

**Dataset S2.** Genomics Dataset. The dataset includes whole genome sequencing of *M. marinum* strains.

